# A Precision Ultrasound-Localized Sonoporation-Equipment (PULSE) Enabling Intratumoral Delivery for Cancer Immunotherapy

**DOI:** 10.1101/2025.07.22.665995

**Authors:** Mengyue Chen, Bohua Zhang, Howuk Kim, Benjamin C. Kreager, Waston Yang, Keren Zhao, Sunho Moon, Huaiyu Wu, Erika J Crosby, Takuya Osada, Jingjie Hu, H. Kim Lyerly, Xiaoning Jiang

## Abstract

Cancer immunotherapy offers a promising long-term treatment for solid tumors by activating the immune system, but side effects and systemic toxicity limit its broad clinical translation. While intratumoral delivery addresses these limitations, elevated stiffness within solid tumors continues to hinder both intracellular and extracellular delivery. Here, a novel therapeutic platform, the Precision Ultrasound-Localized Sonoporation-Equipment (PULSE), is presented to enable spatiotemporally controlled, ultrasound-mediated gene and drug delivery for intratumoral immunotherapy. The PULSE incorporates forward and sideward-looking miniaturized ultrasound transducers, integrated into either a 7-French catheter (cPULSE) or a 16-gauge needle (nPULSE), to minimally invasively sonicate a centimeter-sized tumor. Following the design, prototyping, and characterization of PULSE, its dual capability for intracellular and extracellular delivery is demonstrated through comprehensive *in vitro* cell-based sonoporation tests and phantom-based drug penetration tests. Luciferase assays (activity: ~10^4^ RLU/μg level) in the cell studies confirm significantly enhanced gene transfection with cPULSE, while increased dye diffusion (width: ~1 cm level) in the phantom tests validates improved perfusion with nPULSE. The reported PULSE shows promise for spatiotemporally precise, controlled, and localized therapeutic delivery in early-stage tumors for intratumoral immunotherapy.

## 1. Introduction

Solid tumors are abnormal masses of cells in organs or tissues, which can cause significant morbidity and mortality, threatening human life and burdening healthcare systems.[1,2] Conventional cancer treatments include surgery, radiation therapy, and systemic therapy. However, surgery can be challenging or even infeasible when tumors are located in inaccessible areas or require highly complex and expensive clinical procedures.[3] Radiation therapy can harm normal cells near cancer cells or along the path of the ionizing radiation beam and cause various side effects.[4] Among systemic therapies, both chemotherapy and hormone therapy are struggling with resistance, which undermines their long-term effectiveness.[5,6] Over the past decades, immunotherapy has gained growing interest. It sparks the immune system and thereby obtains durable anti-tumor responses.[7–10] Nevertheless, immunotherapeutic agents can lead to severe side effects, including autoimmunity, nonspecific inflammation, and cytokine storm, which underscores the need for targeted, controlled, and localized delivery.[11,12] A novel delivery strategy, intratumoral injection, has been proposed to limit off-target and systemic toxicity while enhancing immunotherapeutic agent concentrations within solid tumors.[13–16] Despite the burgeoning promise of intratumoral delivery in medical and clinical research, the tumor microenvironment continues to pose a major obstacle to effective drug penetration.[17– 19] Rapid tumor cell proliferation can compress or displace blood and lymphatic vessels, inhibiting drug delivery to cells distal from functional vasculature.[18,19] Immunotherapeutic agents must first penetrate the extracellular matrix within the tumor to reach their target cells and then cross the cell membrane to induce genetic transfection or other gene therapy effects. However, tumor mechanical abnormalities, including 1) elevated solid stresses, 2) elevated interstitial fluid pressure, and 3) increased stiffness, thwart effective drug delivery both extracellularly and intracellularly.[17,18,20,21] Various stimuli-responsive strategies, including light, heat, electricity, magnetism, and ultrasound, have been proposed to enhance both extracellular and intracellular drug delivery in solid tumors while improving the controllability of drug release.[22,23] Among those options, ultrasound is drawing increased attention because it offers finer spatiotemporal resolution and sufficient penetration depth, enabling more precise and effective release than other exogenous stimuli.[24]

Ultrasound, specifically therapeutic ultrasound, uses high-frequency (> 20 kHz) mechanical waves to interact with biological tissues at cellular and molecular levels to achieve therapeutic benefits.[25,26] One of the ultrasound-induced biological effects, sonoporation, has emerged as an effective therapeutic technique for intracellular delivery in cancer immunotherapy.[27–33] With the assistance of ultrasound-responsive cavitation nuclei, including microbubbles, nanodroplets, or microgels, ultrasound waves can enhance cell membrane permeability transiently and increase the uptake of drugs or genes in cells.[34–38] Alongside advances in cavitation-induced sonoporation in the past decades, the underlying mechanisms were gradually revealed, even though not fully understood. As the most broadly used cavitation nuclei, gas-filled, phospholipid-coated microbubbles can expand and shrink with the rarefaction and compression of ultrasound waves.[37] Such a process is called stable cavitation and may occur at lower acoustic pressure conditions. In addition, higher acoustic pressure contributes to an intense radial oscillation and leads to the implosion of microbubbles, enabling inertial cavitation. Both stably and inertially cavitating microbubbles can influence the cell mechanically. Once the cavitating microbubble approaches the cell closely, i.e., usually less than three-quarters of the bubble radius, it may pull and push the cell membrane by microstreaming, jetting, shear stress, or radiation force, to perforate the cell membrane and disassemble the cytoskeleton transiently.[28,36,39,40] Cavitation may also modulate mechanosensitive ion channels to provoke hyperpolarization, further facilitating endocytosis.[28,30,36,41] Through sonoporation, macromolecules such as drugs or genes can be efficiently transported into target cells, aiding in tumor treatment and eradication.[27,31,32] Beyond the cell membrane, the extracellular matrix is another barrier to effective drug delivery in solid tumors.[42–44] Comprising collagens, elastin, and glycoproteins, the extracellular matrix forms a crosslinked network that surrounds and supports cells.[43] In solid tumors, however, activated cancer-associated fibroblasts and high interstitial pressure lead to a stiffer, denser, and more disorganized and crosslinked extracellular matrix, thereby hindering drug penetration.[43,45,46] Recent studies have demonstrated that ultrasound can also enhance extracellular delivery under various sonication conditions.[47,48] High-intensity ultrasound can mechanically or thermally disrupt the tumor’s extracellular matrix, enhancing drug penetration or immune cell infiltration.[49–51] Meanwhile, low-intensity ultrasound can employ acoustic streaming to transport drugs across the matrix crosslinking.[47,52]

The promising potential of ultrasound-mediated intracellular/extracellular delivery attracts considerable interest, fueling ongoing efforts to enhance its delivery efficacy for cancer immunotherapy. Ultrasound-related research efforts have focused on optimizing cavitation nuclei through composition and size adjustments, alongside advancing ultrasound device design via structural and material innovations. For instance, liposomal bubbles encapsulating perfluoropropane for prolonged circulation stability, cationic microbubbles for enhanced electrostatic adhesion to cell membranes, magnetic microbubbles for precise magneto-acoustic modulation, large microbubbles with low ultrasound frequency for intensified cavitation, and smaller bubbles for deeper tumor penetration are employed to boost therapeutic efficacy.[53–58] Yet, current clinical practice still relies on bulky transducers. Encouragingly, microfabrication is propelling miniaturized devices for broader drug delivery, even if some of them are not tailored for tumor treatment. For example, an ultrasound micro-transducer array for site-specific sonoporation, an acoustofluidic platform for high-throughput gene editing, a conformable ultrasound patch for transdermal cosmeceutical delivery, and a wearable ultrasound transducer array for continuous sonodynamic therapy have been successively proposed.[59–62] Nevertheless, both novel miniaturized ultrasound devices and conventional bulky transducers operate extracorporeally, limiting their ability to effectively sonicate tumors behind bones and adipose tissue due to ultrasound attenuation and absorption. Moreover, they cannot precisely and simultaneously co-deliver cavitation nuclei and therapeutic agents to the sonication area. Drawing inspiration from therapeutic intravascular ultrasound transducers, a minimally invasive platform is needed to overcome these limitations and enable spatiotemporally localized ultrasound-mediated gene and drug delivery for explicitly tailored intratumoral immunotherapy.[63–67]

In this study, we report a Precision Ultrasound-Localized Sonoporation-Equipment (PULSE), which is a novel therapeutic platform tailored for spatiotemporally controlled, ultrasound-mediated gene and drug delivery in intratumoral immunotherapy (Fig. 1a). The PULSE comes in two versions: the cPULSE, which is integrated into a catheter, and the nPULSE, embedded within a needle (Fig. 1b). The cPULSE features a single forward-looking stacked transducer, while the nPULSE incorporates three side-looking stacked transducers and one forward-looking stacked transducer arranged in an L-shaped configuration. We first develop the PULSE by designing the device with acoustic-piezoelectric coupled simulations and fabricating it through microfabrication techniques. The developed PULSE is subsequently characterized using electrical and acoustic measurements, while microbubble cavitation is analyzed by theoretical simulations (Fig. 2). To demonstrate intracellular delivery, the nPULSE is used in cell tests to evaluate microbubble cavitation-mediated sonoporation efficiency via luciferase expression (Fig. 3). For extracellular delivery, the cPULSE is applied in phantom tests, where acoustic streaming-driven drug penetration is observed by dye diffusion (Fig. 4). With future *in vivo* studies in animal models and human clinical trials, we envision that the PULSE, as a novel therapeutic platform, can be promising for spatiotemporally precise, controlled, and localized therapeutic delivery in intratumoral immunotherapy.

**Fig. 1.**
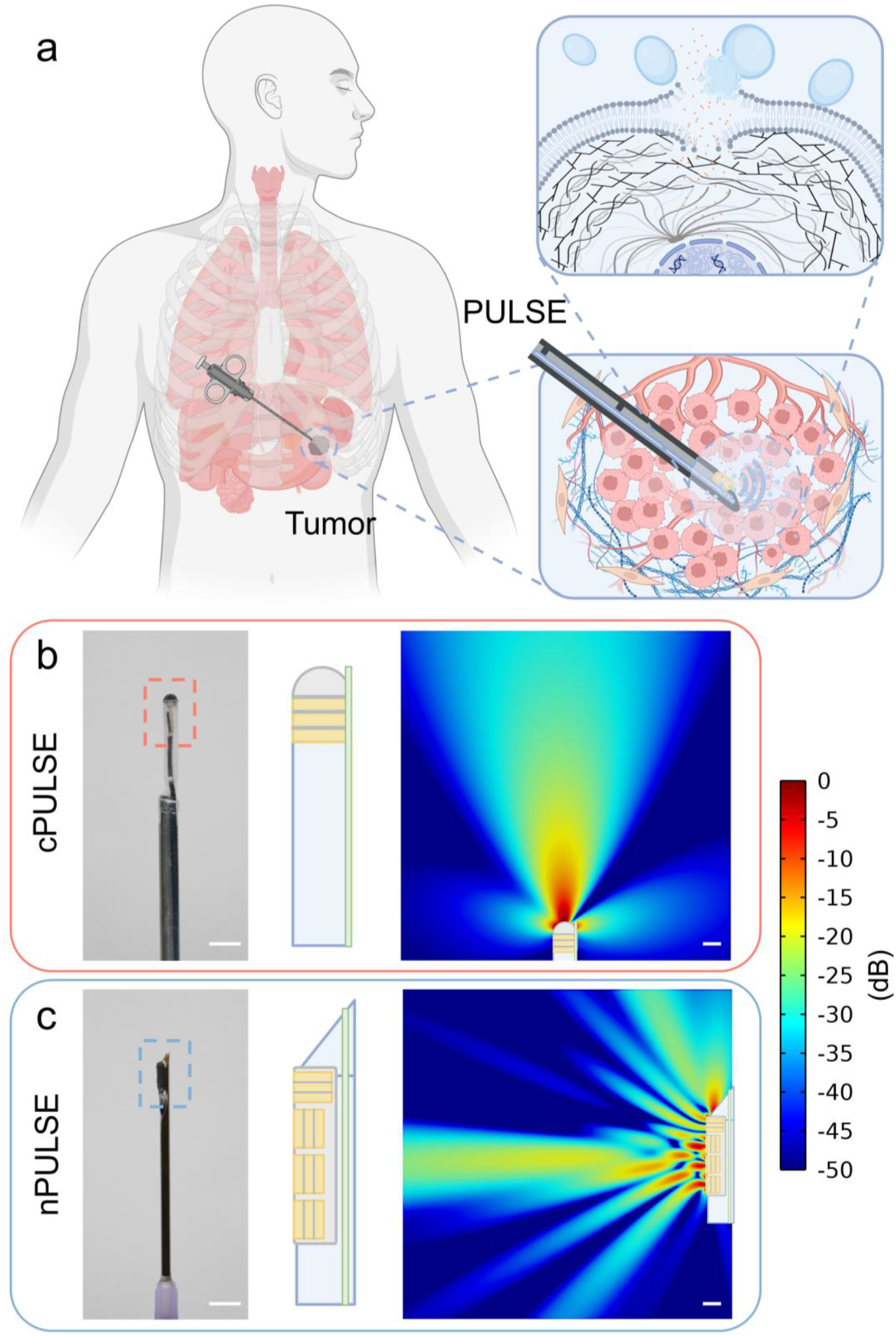
Overview of PULSE. a) Schematic illustration of the PULSE for intratumoral immunotherapy. The minimally invasive PULSE is inserted into the solid tumor to co-deliver both immunotherapeutic agents and cavitation nuclei. By harnessing acoustic streaming and cavitation-mediated sonoporation, immunotherapeutic agents are delivered to target cells both extracellularly and intracellularly, enhancing treatment precision and efficacy. b-c) Photo (left, scale bar: 5 mm), sketch (middle), and simulated acoustic pressure field (right, scale bar: 1 mm) of cPULSE and nPULSE. b) The cPULSE features a single 1.0 MHz PZT-5A stack with a convex matching layer, integrated into a 7-French catheter. c) The cPULSE incorporates four 1.4 MHz PZT-5A stacks arranged in a 3+1 L-shaped configuration, housed within a 16-gauge needle.

**Fig. 2.**
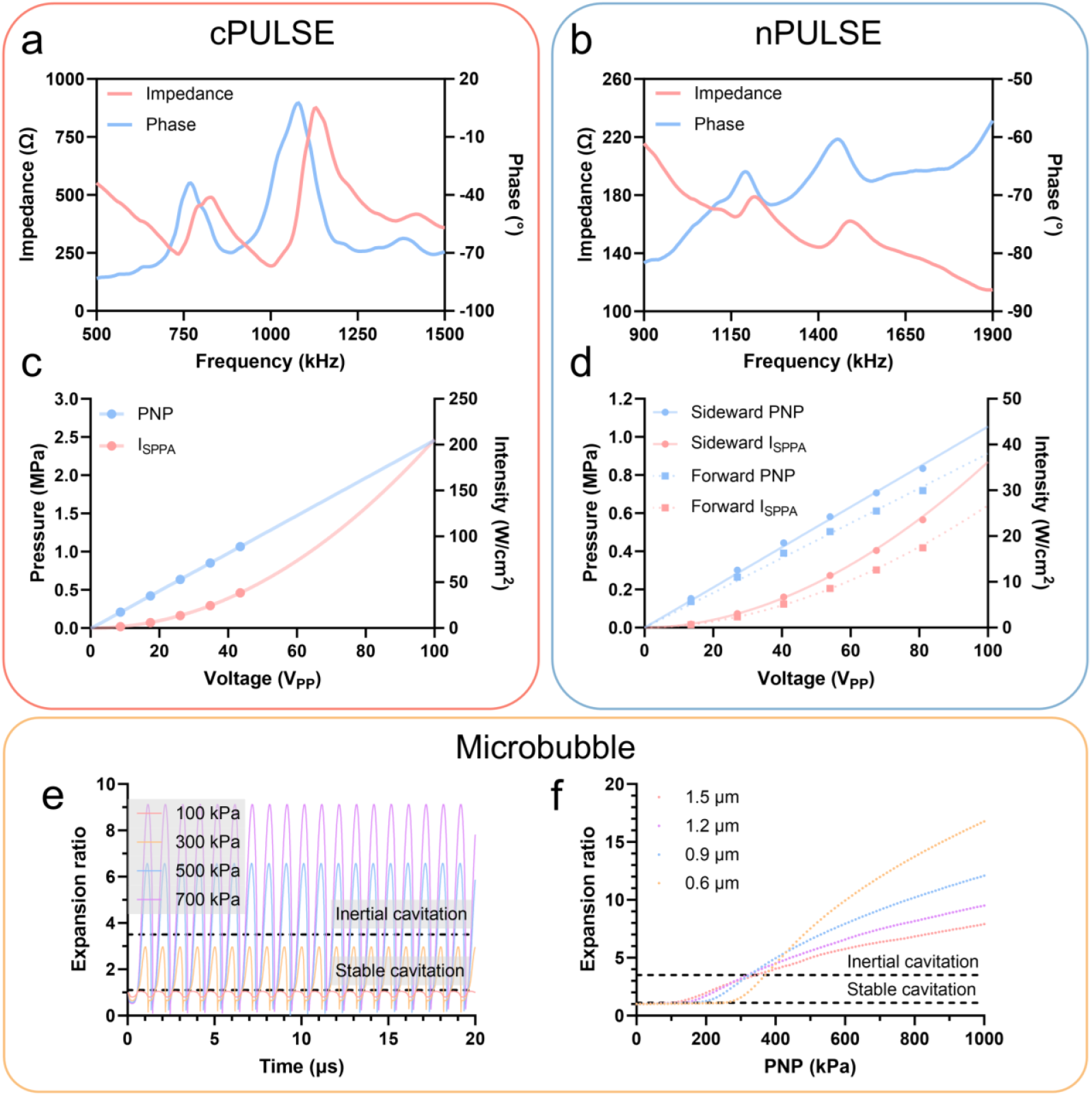
Electrical and acoustic characterization of PULSE and microbubble-induced cavitation analysis. a-d) Measured electrical impedance spectrum (a, b) and acoustic pressure/intensity output (c, d) of cPULSE and nPULSE. The minimum impedance frequency is 1.0 MHz for cPULSE and 1.4 MHz for nPULSE. Both systems can generate the required PNP (0-1 MPa) to induce microbubble-mediated cavitation and acoustic streaming. e-f) Simulated radial oscillations of microbubbles with different diameters (e: 0.9 μm; f: 0.6-1.5 μm) using the Marmottant model, illustrating PNP-dependent shifts between stable and inertial cavitation thresholds.

**Fig. 3.**
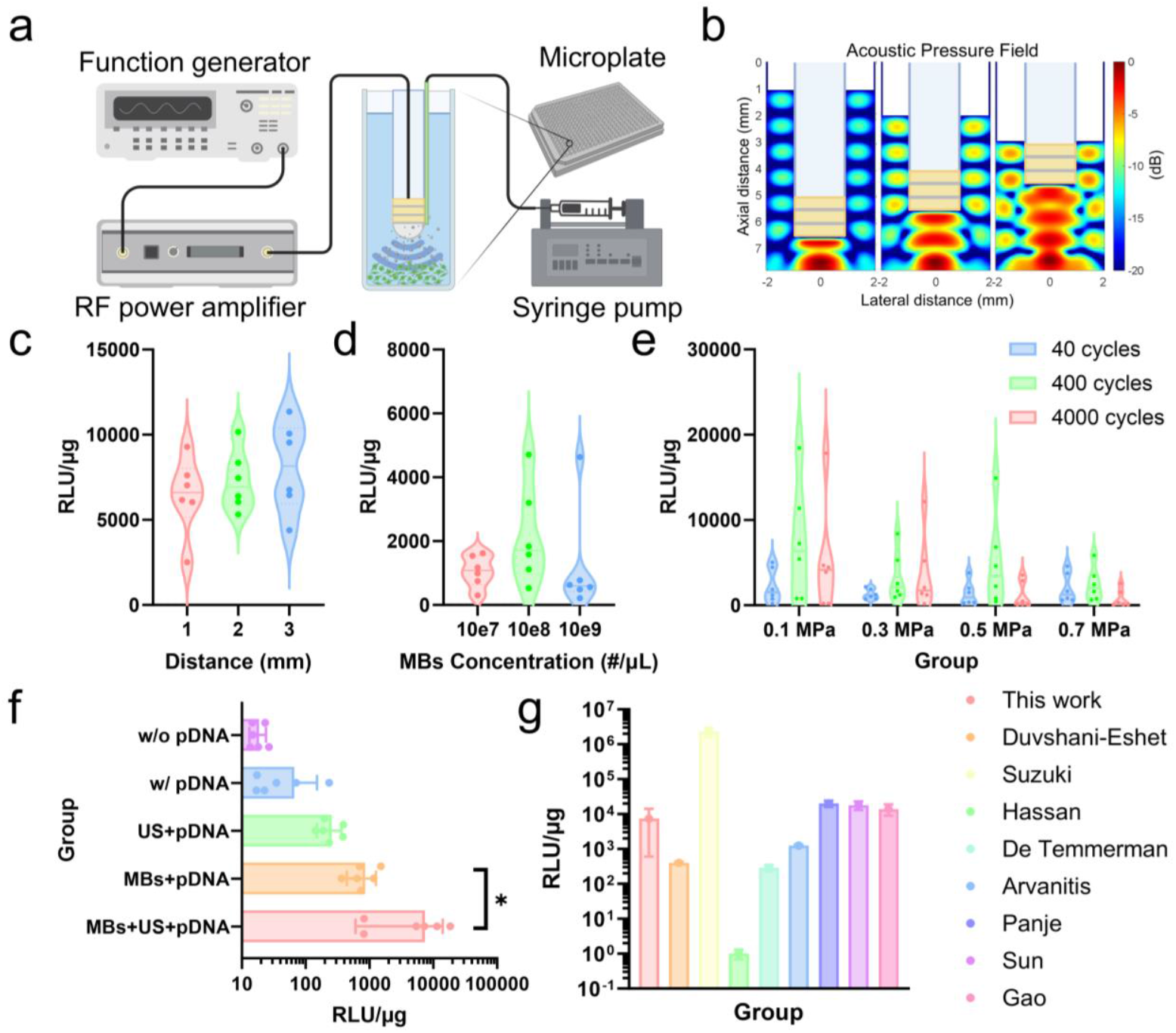
Sonoporation-induced gene delivery. a) Experiment setup for the *in vitro* cell tests. b) Simulated acoustic pressure distribution inside the microplate, demonstrating the impact of standing waves and transducer positioning. c-g) Luciferase activity results from sonoporation tests (n = 6), reflecting the efficacy of intracellular delivery. c) The dependence of gene delivery efficacy on cPULSE positioning, with a distance between cPULSE and microplate varying from 1 mm to 3 mm. d) The dependence of gene delivery efficacy on microbubble concentration, with concentration ranging from 10^7^ to 10^9^ per µL. e) The dependence of gene delivery efficacy on PNP and CN, with PNP varying from 0.1 MPa to 0.7 MPa and CN ranging from 40 to 4000 cycles. f) The comparison between the experiment group (distance: 3 mm; microbubble concentration: 10^7^ #/µL; PNP: 0.1 MPa; CN: 400 cycles) and four control groups, highlighting the significance of sonoporation-induced gene delivery (*p = 0.04). g) The comparison among the experiment results (distance: 3 mm; microbubble concentration: 10^7^ #/µL; PNP: 0.1 MPa; CN: 400 cycles) and several published results, indicating satisfactory sonoporation efficacy.

**Fig. 4.**
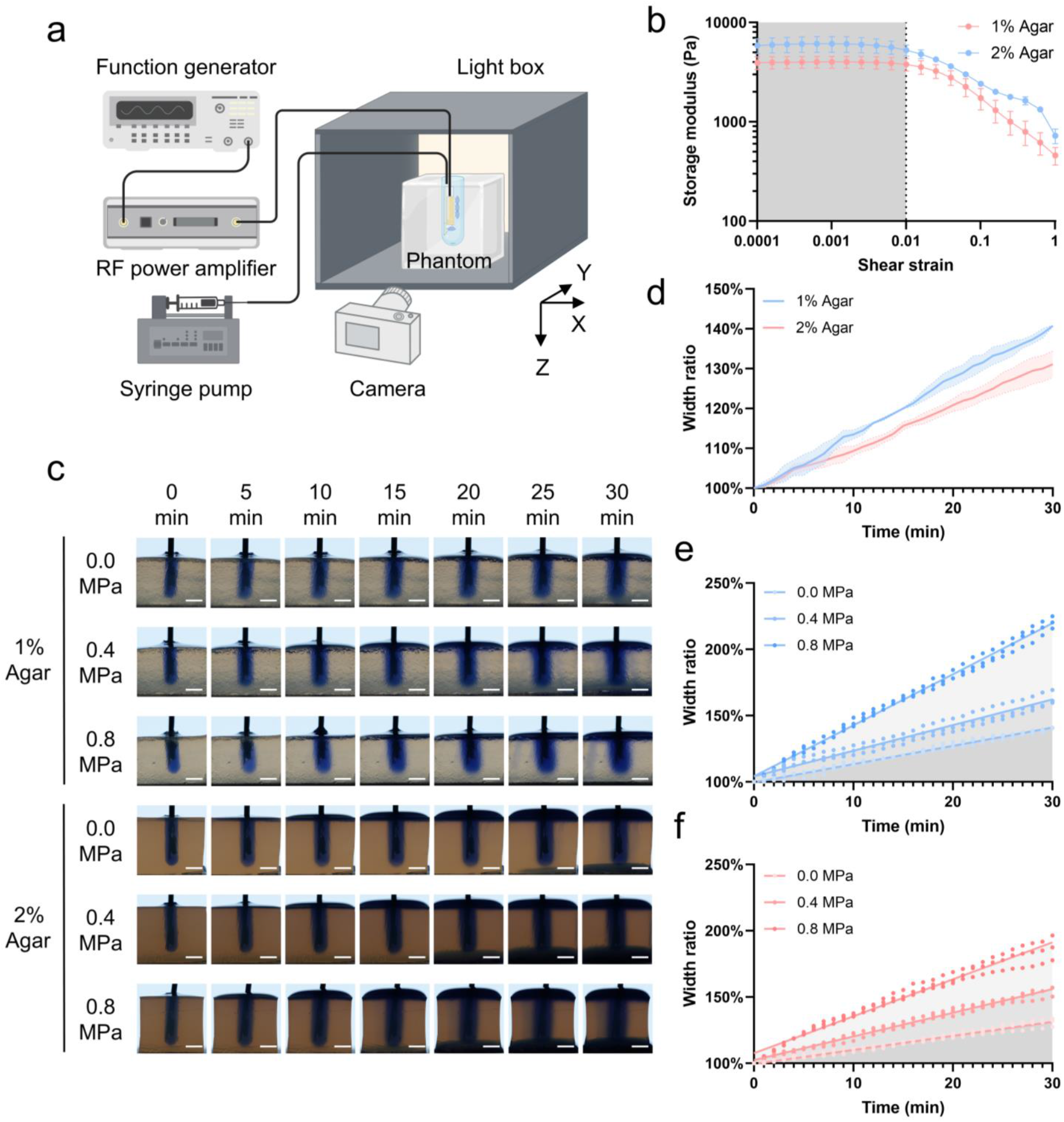
Ultrasound-enhanced drug penetration. a) Experiment setup for the *in vitro* phantom tests. b) Measured shear storage modulus (G’) of 1% and 2% agar phantom (n = 3). c) Representative images illustrating the process of acoustic streaming-enhanced dye diffusion in 1% and 2% agar phantoms with varying PNPs (scale bar: 5 mm). d) The comparison between passive dye diffusion (i.e., control groups) in 1% and 2% agar phantoms indicates that the stiffer, less porous structure of the gel phantom impedes drug penetration. e-f) Measured width changes of the dye diffusion area for demonstrating drug penetration along the x-axis in 1% (e) and 2% (f) agar phantom (n = 3).

## 2. Results

### 2.1. PULSE development

In solid tumors, volumes exceeding 10 mm^3^ are expected to exhibit maximum interstitial fluid pressure, resulting in the minimum achievable drug concentrations within the tumor microenvironment.[68] Given this, we develop PULSE with dual functionality: 1) a millimeter-scale cross-section enabling minimally invasive intratumoral insertion, and 2) the capacity to generate therapeutic effects across centimeter-scale tumor volumes. To achieve minimally invasive insertion, ultrasound transducers are integrated into two distinct devices: a 7-French catheter (2.33 mm outer diameter) for the cPULSE system and a 16-gauge needle (1.65 mm outer diameter) for the nPULSE system, as illustrated in Fig. 1b. To enlarge the treatment area, the ultrasound transducer of cPULSE features a convex matching layer, providing a wider focus compared to a flat matching layer (Fig. S1). Meanwhile, the nPULSE employs ultrasound transducers arranged in a 3+1, L-shaped configuration to cover both forward and sideward zones (Fig. S2). We adopt a stacked design for the ultrasound transducer structure to achieve higher acoustic output pressure at equivalent input voltages, leveraging reduced electrical impedance within identical aperture dimensions. PZT-5A is chosen as the active material to fabricate a three-layer stack, as it strikes a balance between sensitivity and stability. An Al_2_O_3_/epoxy composite matching layer, with an acoustic impedance of ~6 MRayls, is used to optimize acoustic energy transfer from the high-impedance PZT-5A stack (~33 MRayls) to low-impedance biological tissue (~1.5 MRayls). An air-microbubble/epoxy backing layer, with an acoustic impedance of ~500 Rayls, is used to suppress backward-propagating ultrasound waves and enhance forward transmission of acoustic energy. For the design of dimensional parameters, we determine the thickness of each PZT layer by referencing the ultrasound frequencies used in published *in vitro* sonoporation studies (Fig. S3). In the cPULSE, the PZT layer thickness is set to 300 µm to achieve a 1.0 MHz resonant frequency, following the half-wavelength-thick active layer design principle. In contrast, the PZT layer thickness in nPULSE is reduced to 200 µm to accommodate space limitations within a 16-gauge needle, resulting in a resonant frequency of 1.4 MHz. The cPULSE utilizes a PZT stack with a 1.6 × 1.6 mm^2^ aperture, while the nPULSE employs PZT stacks featuring a reduced 1.0 × 1.0 mm^2^ aperture. The designed cPULSE and nPULSE are fabricated through microfabrication techniques, including lapping, bonding, and dicing (see Experimental Section).

### 2.2. PULSE characterization

The electrical and acoustic performance of the prototyped PULSE is assessed via impedance response characterization and acoustic pressure output measurement. As illustrated in Fig. 2, the resonant frequency of cPULSE in thickness mode is 1.0 MHz, whereas nPULSE exhibits a higher resonant frequency of 1.4 MHz due to its thinner stack. The electrical impedances at the resonant frequency for cPULSE and nPULSE are 193.6 Ω and 144.2 Ω, respectively. At 1 kHz, cPULSE has a dielectric loss of 12.2 mU and a capacitance of 540.9 pF, while nPULSE has a dielectric loss of 11.5 mU and a capacitance of 781.9 pF. The measured acoustic pressure (i.e., peak negative pressure, PNP) and intensity (i.e., spatial-peak pulse-average intensity, I_SPPA_) output for cPULSE and nPULSE at their respective natural focal points are displayed in Fig. 2. The cPULSE exhibits a transmission sensitivity of 24.6 kPa/V_PP_, whereas the nPULSE demonstrates sensitivities of 10.1 kPa/V_PP_ for three side-looking elements and 8.6 kPa/V_PP_ for one forward-looking element. These transmission sensitivities enable the PULSE to achieve the target acoustic pressure range (<1 MPa) required for sonoporation while operating within practical driving voltages (<100 V_PP_), avoiding material depolarization or safety concerns for intracorporeal applications.

We also predict stable and inertial cavitation through theoretical simulation by calculating the microbubble expansion ratio. Fig. 2e presents the radial oscillation dynamics of a 0.9 μm diameter microbubble under 1.0 MHz ultrasound exposure at varying PNP amplitudes. At a PNP of 0.1 MPa, the microbubble exhibits minimal oscillations with a sinusoidal waveform and achieves an expansion ratio of ~1.1, marking the threshold for stable cavitation.[69] At a PNP of 0.3 MPa, microbubble oscillations transition to a non-sinusoidal waveform, indicating more intense cavitation. At a PNP higher than 0.5 MPa, microbubble expansion ratios exceed a critical threshold of 3.5, showing the transition from stable to inertial cavitation.[69] According to the microbubble size distribution (Fig. S4), four representative diameters (from 0.6 to 1.5 μm range) are selected, and their oscillation dynamics are further investigated under varying PNPs. The results, shown in Fig. 2f, indicate that under 1.0 MHz sonication, stable cavitation occurs at PNPs between 0.1 and 0.4 MPa, while PNPs greater than 0.4 MPa induce inertial cavitation.

### 2.3. Sonoporation-induced gene delivery

To demonstrate intracellular delivery through sonoporation, we employ the cPULSE to sonicate HEK-293 cells in a microplate containing plasmid DNA and microbubbles (Fig. 3a). Given that ultrasound reflection may affect the *in-situ* acoustic pressure level, the acoustic pressure distribution within a microplate well is simulated. Fig. 3b shows the simulated propagation of the 1.0 MHz ultrasound wave from the cPULSE surface (positioned at varying heights) toward the bottom of the microplate well. The acoustic impedance mismatch between water and the polystyrene microplate induces ultrasound wave reflection at the bottom, while the water-air interface produces a secondary reflection at the top. The dual reflections generate standing waves within the microplate well cavity, and their pressure amplitude increases as the transducer is positioned closer to the top. Additionally, a wider and higher acoustic pressure field develops near the microplate well bottom, suggesting its suitability for sonicating cells. Compared to the free-water cPULSE configuration, positioning the PULSE at heights of 1 mm, 2 mm, and 3 mm in the microplate results in acoustic pressure increases of 17.5%, 63.2%, and 48.8%, respectively. To further investigate the impact of transducer height on sonoporation efficiency, we conduct *in vitro* experiments. The results (Fig. 3c) suggest that elevating the transducer height could enhance sonoporation efficacy. This could be attributed to the larger high-pressure area generated at the microplate bottom when the transducer height is 3 mm, which may sonicate a greater number of cells (Fig. 3b). We proceed to study the impact of microbubble concentration and find that a concentration of 10^8^/µL achieves higher sonoporation efficacy than both 10^7^/µL and 10^9^/µL (Fig. 3d).

After determining the optimal microbubble concentration (i.e., 10^8^/µL) and transducer heights (i.e., 3 mm), we systematically assessed sonication variables such as PNP and cycle number (CN) to evaluate their influence on sonoporation efficiency. PNP of 0.1, 0.3, 0.5, and 0.7 MPa, and CN of 40, 400, and 4000 cycles are tested. The results illustrated in Fig. 3e suggest that excessively high PNP (i.e., 0.7 MPa) and insufficient CN (i.e., 40 cycles) are detrimental to improving sonoporation efficiency. This may result from increased cell death caused by irreversible cell membrane perforation under excessive inertial cavitation, and reduced microbubble-cell contact probability during sonication under shorter pulse durations.[36] Notably, under the CN condition of 400 cycles, luciferase activity exhibits dual peaks at 0.1 MPa and 0.5 MPa, which may be attributed to stable and inertial cavitation contributions, respectively.[70] At 4000 cycles, lower PNP results in higher luciferase activity, suggesting that stable cavitation under increased cycle numbers prolongs effective sonication duration, thereby improving sonoporation efficiency. Although inertial cavitation enhances sonoporation efficiency by generating larger pores in the cell membrane, its efficacy diminishes at higher CN due to abrupt microbubble collapse. To further evaluate intracellular delivery efficacy via cPULSE, the experimental group (PNP: 0.1 MPa, CN: 400 cycles) is benchmarked against control groups (Fig. 3f) and existing literature data (Fig. 3g) [53,55,71–76]. Given the lower luciferase activity range observed in the control groups, the y-axis scale is adjusted from linear to logarithmic to enhance data visibility. The results show that ultrasound combined with microbubbles significantly enhances gene transfection, achieving efficiency comparable to existing literature, which suggests cPULSE as a promising platform for sonoporation-induced gene delivery.

### 2.4. Ultrasound-enhanced drug penetration

To demonstrate extracellular delivery via acoustic streaming, we utilize the nPULSE to sonicate tumor-mimicking gel phantoms containing a dye solution within a customized photo studio light box (Fig. 4a). The dye serves as a drug surrogate, and drug penetration is visualized by tracking its diffusion to assess spatial permeation. Agar phantoms of varying concentrations are employed to mimic tumors. To assess their ability to replicate the mechanical properties of tumors, we measure their shear modulus as this parameter reflects the crosslinking density within a tumor’s extracellular matrix. Fig. 4b presents the measured shear modulus (G’) for agar phantoms with 1% and 2% concentrations. The mean ± standard deviation values for shear strain < 0.01 are 3952 ± 66.4 Pa and 5904 ± 259.2 Pa, respectively, which are higher than the reported value of human pancreatic tissue (2800 ± 600 Pa).[77] This suggests that agar phantoms are effective models for mimicking tumor tissue.[78] Additionally, the measured shear loss modulus (G’’) in Fig. S5 indicates that the 1% agar phantom exhibits more liquid-like behavior than the 2% agar phantoms under deformation-limited conditions (shear strain < 0.01).

After confirming the suitability of agar phantoms, we subject them to sonication with PNPs ranging from 0.0 to 0.8 MPa. Notably, unlike our previous studies that employed microbubbles to amplify ultrasound effects, here we exclude microbubbles to assess how acoustic streaming, in the absence of cavitation, enhances drug penetration.[79] Fig. 4c illustrates the dye diffusion process over time in 1% and 2% agar phantoms under different PNPs, demonstrating that the dye spreads over a larger area at higher PNP. To quantify the dye diffusion process, we analyze the dye’s width (x-axis), depth (z-axis), and color intensity (y-axis) to evaluate its penetration across three-dimensional space (x, y, z). Since the initial conditions of each experimental group may vary slightly, changes in width (x-axis), depth (z-axis), and color intensity (y-axis) are normalized by expressing each parameter as a ratio of its value at the nth minute to its baseline (0-minute) value (see Experimental Section).

Fig. 4d depicts the x-axis dye diffusion comparison between 1% and 2% agar phantoms under zero PNP conditions (i.e., control group). As anticipated, higher agar concentration correlates with reduced drug penetration, aligning with the elevated shear storage modulus (Fig. 4b) and darker color intensity (Fig. 4c) observed in these samples. Fig. 4e and Fig. 4f show the x-axis dye diffusion profiles under various PNP conditions in 1% and 2% agar phantoms, respectively. Drug diffusion exhibits a magnitude-dependent enhancement under applied PNP, with 0.4 MPa increasing diffusion by ~20% relative to the 0.0 MPa baseline in both 1% and 2% agar phantoms. At 0.8 MPa PNP, diffusion enhancement reaches ~80% in the 1% agar phantom but decreases to ~55% in the 2% agar phantom, indicating that the elevated crosslinking density inherent to higher-concentration agar imposes a structural limitation on acoustic streaming-driven drug diffusion, even at high acoustic pressure magnitude. Similar phenomena are also observed in the dye diffusion profiles along the y-and z-axes, as shown in Fig. S6 and Fig. S7.

## 3. Discussion

Bulky, extracorporeal ultrasound devices are the prevailing approach used in medical and clinical studies for ultrasound-mediated drug and gene delivery in cancer immunotherapy. However, these devices struggle to effectively treat tumors located behind bony structures or within fatty tissue, and they face challenges in achieving precise spatiotemporal delivery of acoustic energy, cavitation nuclei, and therapeutic agents to target sites. In this work, we present PULSE, a novel, miniaturized, and minimally invasive ultrasound platform designed to merge the advantages of intratumoral injection with sonoporation-mediated immunotherapy, enabling spatiotemporally precise, controlled, localized, and effective therapeutic delivery for cancer treatment. Two unique versions of PULSE, cPULSE and nPULSE, are proposed, developed, and characterized. The cPULSE is designed for implantation into tumors following minimally invasive surgery, whereas the nPULSE enables direct percutaneous insertion into the tumor. To expand and deepen the sonication area, the forwarding element in cPULSE is integrated with a convex matching layer, while the three side-looking elements in nPULSE are aligned in a linear arrangement. Fig. 1b and Fig. 1c demonstrate that the simulated ultrasound focal zone (−12 dB) achieves centimeter-scale dimensions, sufficient to sonicate early-stage tumors such as T1 and T2, which are typically < 5 cm in size.

Then, the cPULSE successfully demonstrates intracellular delivery via sonoporation, as validated through *in vitro* cell tests. *In vitro* experimental parameters, such as transducer-to-cell distance and microbubble concentration, have been demonstrated to influence gene transfection efficiency (Fig. 3b-3d). These findings confirm that optimizing microbubble-cell contact probability within the effective sonication field is critical for maximizing intracellular delivery efficacy. Additionally, sonication parameters, including PNP and CN, are pivotal in modulating the intensity of stable or inertial cavitation, thereby directly enhancing sonoporation success. Stable cavitation is optimized under conditions of low PNP and high CN, whereas inertial cavitation necessitates high PNP but shows no enhancement from elevated CN (Fig. 2c and Fig. 3e). Notably, *in situ* PNP, rather than free-water PNP, is the critical factor for precisely and effectively controlling cavitation. However, measuring the *in situ* PNP using hydrophones is challenging due to spatial and structural constraints. Conventional extracorporeal ultrasound devices need to account for acoustic attenuation or distortion, whether caused by the ultrasound wave traveling through tissue during *in vivo* studies or through plastic containers such as microplates, Petri dishes, and centrifuge tubes during *in vitro* tests. Minimally invasive intracorporeal ultrasound devices are less affected by these issues during *in vivo* studies, as the treatment area is close to the transducer surface. However, *in vitro* testing may amplify *in situ* pressure due to acoustic reflections and standing wave formation (Fig. 3b). Overall, our findings indicate that an *in situ* PNP less than 0.5 MPa is effective for triggering sonoporation under 1.0 MHz sonication.

Lastly, the nPULSE successfully demonstrates extracellular delivery via acoustic streaming, as validated through *in vitro* phantom tests. By changing its concentration, the agar phantom is shown to effectively mimic not only normal tissue but also tumor tissue (Fig. 4b).[80] As shown in Fig. 4c, the dye diffusion process reveals that precise control of PNP and sonication time enables tunable regulation of the dye’s spatial distribution. Following 30 minutes of 0.8 MPa PNP sonication, the dye expanded from an initial diameter of 4.0 mm (i.e., the diameter of the pre-drilled hole) to approximately 8.8 mm in 1% agar and to 7.5 mm in 2% agar. Notably, the nPULSE is inserted into the phantom’s pre-drilled hole to avoid crack formation during insertion. We acknowledge that this approach increases the contact area between the liquid dye and the solid phantom, thereby magnifying dye diffusion. To address this limitation, we propose integrating multi-port injection channels into the next-generation design of PULSE. Also, in contrast to our earlier findings, this study demonstrates that acoustic streaming alone, without cavitation, enhances dye diffusion, offering greater flexibility in optimizing injection and sonication sequencing strategies for future *in vivo* applications. We observe that higher-concentration agar imposes structural constraints on acoustic streaming-driven drug diffusion (Fig. 4e and Fig. 4f). This finding suggests that physically disrupting the crosslinking structure of tumor-mimicking phantoms or tumor tissue, for example, via histotripsy, could improve drug delivery efficacy.[49]

This work represents an important step forward in the development of intratumoral ultrasound devices for minimally invasive cancer immunotherapy. Through comprehensive *in vitro* testing, it lays the groundwork for future *in vivo* animal studies and clinical translation. Future work will focus on: 1) engineering a next-generation PULSE system with circumferential side-looking transducer arrays and multi-port injection channels to enhance therapeutic coverage for advanced-stage tumors (e.g., T3/T4); and 2) performing animal trials in rat or rabbit models using optimized injection-sonication sequence protocols to evaluate cancer treatment efficacy. With this, we envision PULSE as a promising and powerful alternative for achieving spatiotemporally precise, controlled, localized, and effective intratumoral immunotherapy.

## 4. Methods

### Device Fabrication

The PULSE was fabricated following our previously described multi-layer, stacked transducer procedure.[65–67,81] Using a lapper (PM5, Logitech Ltd., UK), we first reduced the thickness of the commercial PZT 5A plate (SM412, Steiner & Martins Inc., USA) to 300 µm for the 1.0 MHz cPULSE and 200 µm for the 1.4 MHz nPULSE. Thermionics e-beam evaporation was then used to deposit 200/10 nm Au/Ti electrodes on both the top and bottom surfaces of the PZT plates. Three PZT 5A plates were stacked using steel-reinforced epoxy (8265S, J-B Weld Company, USA) under controlled pressure via a custom-designed jig, achieving a uniform bond layer thickness of approximately 20 µm. The matching layer was fabricated by mixing and centrifuging 50 nm Al_2_O_3_ particles (ALR-1005-01, Pace Technologies, USA) with epoxy resin (EPO-TEK 301, Epoxy Tech. Inc., USA) at a 4:1 weight ratio, and then lapped to a thickness of 1/4 wavelength after curing the mixture. Notably, the matching layer of the cPULSE was cured into a convex shape using a custom concave mold to enlarge the focal area. The backing layer was fabricated by mixing and centrifuging air microbubble powders (Blatek Inc., USA) with epoxy resin (EPO-TEK 301, Epoxy Tech. Inc., USA) at a 3:1 weight ratio, and then lapped to a thickness of 5 wavelengths after curing the mixture. Subsequently, the matching and backing layers were bonded to the respective front and rear surfaces of the PZT stack, using steel-reinforced epoxy as the bonding agent. After curing at 40 °C for 24 hours, the entire stack was precisely sliced into the desired aperture size using a dicing saw (DAD322, DISCO Inc., Japan). Specifically, the cPULSE element has an aperture of 1.6 × 1.6 mm^2^, while the nPULSE element has an aperture of 1.0 × 1.0 mm^2^. The three side-looking stacks of the nPULSE were partially sliced to retain their structural connection through the backing layer and mounted to the rear side of the forward-looking stack. Next, a thin Al_2_O_3_/epoxy layer was applied to the stack’s side to isolate unwanted electrodes, while a thin silver epoxy layer (E-Solder 3022, Von-Roll Inc., USA) was used to interconnect necessary electrodes, which were then linked to a coaxial cable (5381-006, AWG 38, Hitachi Cable America Inc., USA). The transducer structure of PULSE was coated with a 10 µm-thick parylene C layer as a passivation layer using a parylene coater (SCS Labcoter, Coating Systems Inc., USA). Finally, we integrated the transducer structure into either a catheter or needle to complete the respective PULSE. For the cPULSE, the forward-looking transducer was integrated into a 2.3 mm outer-diameter catheter, which featured an adjacent 1.0 mm outer-diameter injection lumen. In contrast, the nPULSE housed an L-shaped transducer configuration (three side-looking and one forward-looking stacks) within a 16-gauge needle, along with a 1.0 mm outer-diameter injection lumen.

### Device characterization

We characterized the prototype PULSE both electrically and acoustically. First, an impedance analyzer (4294A, Agilent Technologies Inc., USA) was used to measure the electrical impedance and phase spectrum over the relevant frequency range, as well as the dielectric loss and capacitance at 1 kHz. Then, a calibrated bullet-type hydrophone (HGL-0085, ONDA Corporation, USA) paired with a 20 dB preamplifier (AH-2020, ONDA Corporation, USA) was used to measure the acoustic pressure output. As shown in Fig. S8, a water tank containing degassed and deionized water housed the PULSE and hydrophone. A 10-cycle sinusoidal pulse within a 1-ms signal was generated by a function generator (Agilent 33250A, USA), amplified by an RF (radio frequency) power amplifier (Amplifier Research 75A250A, USA), and transmitted to drive the PULSE. The acoustic pressure signal was recorded across varying input voltages using a digital oscilloscope (DSO7104B, Agilent Technologies Inc., USA), and the hydrophone was positioned at the focal point using a 3-axis positioning system. The forward-looking element’s focal point lies approximately 1.0 mm from the transducer surface, whereas the side-looking element’s focal point is about 1.5 mm away. The transmission sensitivities of PULSE, involving acoustic pressure and intensity, were extrapolated using linear and second-order polynomial trendlines, respectively.

### Numerical simulation

Three distinct simulations were conducted in this study, including an acoustic-piezoelectric coupled simulation, a theoretical microbubble cavitation simulation, and an acoustic simulation. The acoustic-piezoelectric coupled simulation was performed using the finite element method analyzer COMSOL Multiphysics (6.3, COMSOL Inc., Sweden) to assist the design of PULSE. The mechanical and acoustic performance of the PULSE was simulated within a two-dimensional plane, incorporating pressure acoustics (frequency domain), solid mechanics, and electrostatics modules. The detailed material properties of active, matching, and bonding layers are summarized in Table S1. The theoretical microbubble cavitation simulation was performed using the Marmottant model to predict the radial oscillations of microbubbles.[82] The effects of acoustic pressure on microbubble expansion ratio were analyzed by numerically solving the modified Rayleigh-Plesset equation using an ordinary differential equation solver (i.e., ODE45) in MATLAB (R2023b, MathWorks Inc., USA).[83] The detailed microbubble properties and sonication parameters are listed in Table S2. The acoustic simulation was performed using a MATLAB toolbox, k-Wave, to optimize the *in vitro* cell test setup.[84] The ultrasound wave propagation of the cPULSE in a constrained area (i.e., polystyrene microplate) was modeled in the time domain. Exploiting the axial symmetry of each microplate well, the computational domain was reduced to a two-dimensional water-filled plane bounded by air and polystyrene structure. The detailed acoustic properties of the polystyrene, water, and air are provided in Table S3.

### Preparation for the cell and phantom tests

The cell test preparation entails preparing cells, DNA, and microbubbles, while the phantom test preparation involves fabricating and characterizing the phantom. For the cell tests, HEK (Human Embryonic Kidney) 293 cell line, plasmid DNA encoding pCDH (protocadherin)-LUC (Luciferase), and commercial microbubbles (VesselVue, SonoVol Inc., USA) were prepared. The cells mixed with PBS (phosphate-buffered saline) (Gibco, Thermo Fisher Scientific, USA) were seeded into a clear 384-well microplate (Falcon, Corning, USA) at 12,000 cells/well (3×10^5^ cells/mL in 40 µL/well), followed by the addition of plasmid DNA (0.4 µg/well at 20 µg/mL in 20 µL/well). The microbubbles consist of a phospholipid shell encapsulating a perfluorobutane core, with a mean diameter of 0.9 ± 0.49 µm. Size distribution data are shown in Fig. S4, provided by the vendor (SonoVol Inc., USA). Before sonication, the microbubbles were activated for 45 s using a shaker (Vialmix, Lantheus Medical Imaging Inc., USA) and then diluted in PBS within a 10 mL syringe (BH Supplies, USA) to achieve a final concentration of 8.7×10^8^ bubbles/mL. For the phantom tests, tumor-mimicking gel phantoms were prepared using agar (A9799, MilliporeSigma, USA) at different powder concentrations (i.e., 1% and 2%, w/v) in PBS. 2 g and 4 g agar powder were individually poured into 200 mL PBS and subsequently heated to temperatures exceeding 100 °C using a magnetic stirring hot plate (PC420D, Corning, USA) under continuous agitation. After complete dissolution, the homogeneous mixture was cast into a customized mold to produce a cubic phantom embedded with a 4.0 mm diameter well. The mixture was cooled gradually to room temperature and then refrigerated to finalize the solidification process. In addition, the shear modulus of 1% and 2% (w/v) agar phantoms was measured at room temperature using a rheometer (MCR 302e, Anton Paar, Austria) with a 25 mm parallel sandblasted plate. Values were extracted from the linear viscoelastic region during frequency sweeps performed at 1 rad/s.[77]

### Procedure for the cell and phantom tests

During cell and phantom tests, both the cPULSE and nPULSE were powered by an RF amplifier and regulated by a function generator. The corresponding sonication parameters are summarized in Table 1. For the cell tests (Fig. 3a), the cPULSE was inserted into the microplate to deliver microbubble solution, with infusion controlled by a syringe pump (NE-1010, New Era Pump Systems, USA) at a rate of 0.1 mL/min. Before sonication, 30 µL of the microbubble solution was infused into each well via injection tube, resulting in a total volume of approximately 90 µL. Using a holder equipped with a 3-axis motion stage, the distance between the cPULSE’s surface and the microplate bottom was precisely set to a specified value (e.g., 2 mm), ensuring no air gap was trapped between the cPULSE and the microplate. After sonication, cells were incubated at 37 °C in a CO_2_ incubator for 24 hours before assessing sonoporation efficiency. For the phantom tests (Fig. 4a), the nPULSE was positioned within the phantom’s pre-drilled hole to prevent cracking during needle insertion. Trypan blue solution (Gibco, Thermo Fisher Scientific, USA) mixed with PBS was injected through the injection lumen, with a syringe pump maintaining a constant flow rate to ensure uniform fluid pressure and a consistent fluid height. Meanwhile, the nPULSE sonicated the solution for 30 minutes while a CMOS (complementary metal-oxide-semiconductor) camera (ILCE-7M4, Sony, Japan) captured the dye diffusion process at 1-minute intervals.

**Table 1.** Key sonication parameters for the cell and phantom tests.

|  |  |  |  |
| --- | --- | --- | --- |
| Cell tests | Frequency<br>1.0 MHz | Pressure<br>0.1 - 0.7 MPa | Mechanical index<br>0.1 - 0.7 |
|  | Cycle number<br>40, 400, 4000 | Pulse repetition period<br>0.5, 5, 50 ms | Sonation duration<br>30 s |
| Phantom tests | Frequency<br>1.4 MHz | Pressure<br>0.4, 0.8 MPa | Mechanical index<br>0.34, 0.68 |
|  | Cycle number<br>280 | Pulse repetition period<br>1 ms | Sonation duration<br>30 mins |

### Experiment assessment and statistical analysis

To assess sonoporation efficiency, cells from each well were harvested and lysed in the lysing buffer. Luciferase activity was quantified by adding luciferin using a luminometer (GloMax, Promega Corporation, USA), with results expressed as relative light units (RLU) per μg protein. To evaluate drug perfusion, changes in dye width, depth, and color intensity were quantified using ImageJ Fiji software (1.54, National Institutes of Health, USA).[85] The corresponding equations used for normalizing dye diffusion along the x-, y-, and z-axes are as follows:

|  |  |  |
| --- | --- | --- |
| x-axis | Width ratio | $Width\ ratio = \frac{width\ at\ nth\ minute}{width\ at\ 0\ minute}$ |
| y-axis | Intensity increase | $Intensity\ increase = 1 - \frac{color\ intensity\ at\ nth\ minute}{color\ intensity\ at\ 0\ minute}$ |
| z-axis | Depth ratio | $Depth\ ratio = \frac{depth\ at\ nth\ minute}{depth\ at\ 0\ minute}$ |

Cell tests were performed in six independent biological replicates (n = 6), and phantom tests were carried out in three independent experimental replicates (n = 3). Test results between control and experimental groups were analyzed using unpaired t-tests, with statistical significance set at p < 0.05.

## Acknowledgments

M.C. thanks Xiangming Xue and Xingru Ma for fruitful discussions on data analysis of phantom tests and concepts of tumor microenvironment. The work was supported by the Duke University Medical Center.

## Appendix A. A. Supplementary data

Supplementary data for this article can be found online.

