## Supplementary Data for "A Precision Ultrasound-Localized Sonoporation-Equipment (PULSE) Enabling Intratumoral Delivery for Cancer Immunotherapy"

**This PDF file includes:**

- supplementary tables S1 to S3
- supplementary figures S1 to S8

**Table S1.** Material properties of the acoustic-piezoelectric coupled simulation

| Properties | Active layer | Properties | Matching layer | Bonding layer |
|---|---|---|---|---|
| Material | PZT-5A | Material | $Al_2O_3$/Epoxy | Epoxy |
| Density | 7750 kg/m$^3$ | Density | 2700 kg/m$^3$ | 1200 kg/m$^3$ |
| $C_{33}^E$ | 111 GPa | Young's modulus | 11.5 GPa | 10 GPa |
| $e_{33}$ | 15.8 C/m$^2$ | Poisson's ratio | 0.32 | 0.35 |

**Table S2**. Sonication parameters and material properties of the theoretical microbubble cavitation simulation

| Properties | Value | Properties | Value |
|---|---|---|---|
| Frequency | 1 MHz | Water density | 998 kg/m$^3$ |
| Sound speed in water | 1489 m/s | Water surface tension | 0.072 N/m |

**Table S3.** Material properties of the acoustic simulation

| Materials | Polystyrene | Water | Air |
|---|---|---|---|
| Sound speed | 2400 m/s | 1489 m/s | 343 m/s |
| Density | 1050 kg/m$^3$ | 998 kg/m$^3$ | 1 kg/m$^3$ |
| Acoustic attenuation | 11.5 GPa | 11.5 GPa | 10 GPa |

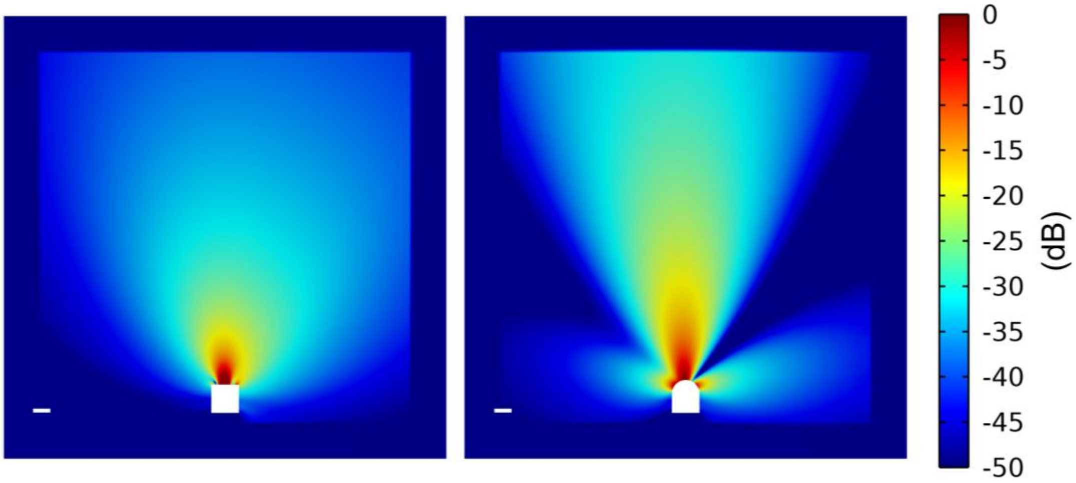


**Fig. S1**. Simulated acoustic pressure field of cPULSE with flat (left) and convex (right) matching layer (scale bar: 1 mm). The convex matching layer contributes to a broader and deeper ultrasound focal area.

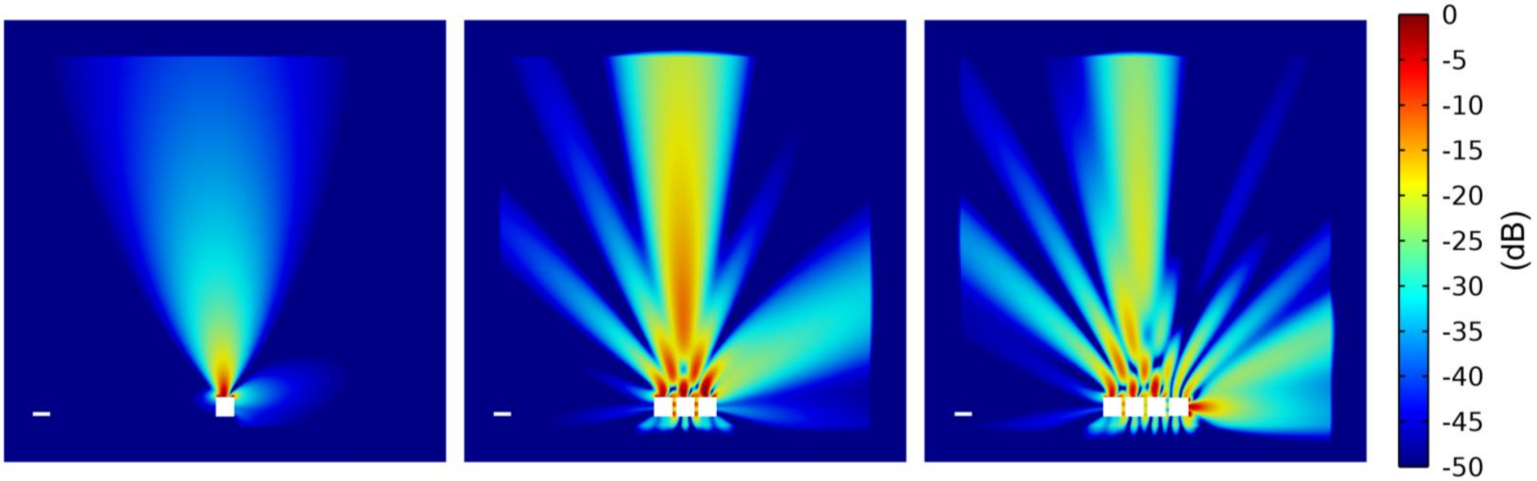

**Fig. S2.** Simulated acoustic pressure field of nPULSE with one, three, and four stacks. Multiple sideward and forward stacks enlarge the ultrasound focal area.

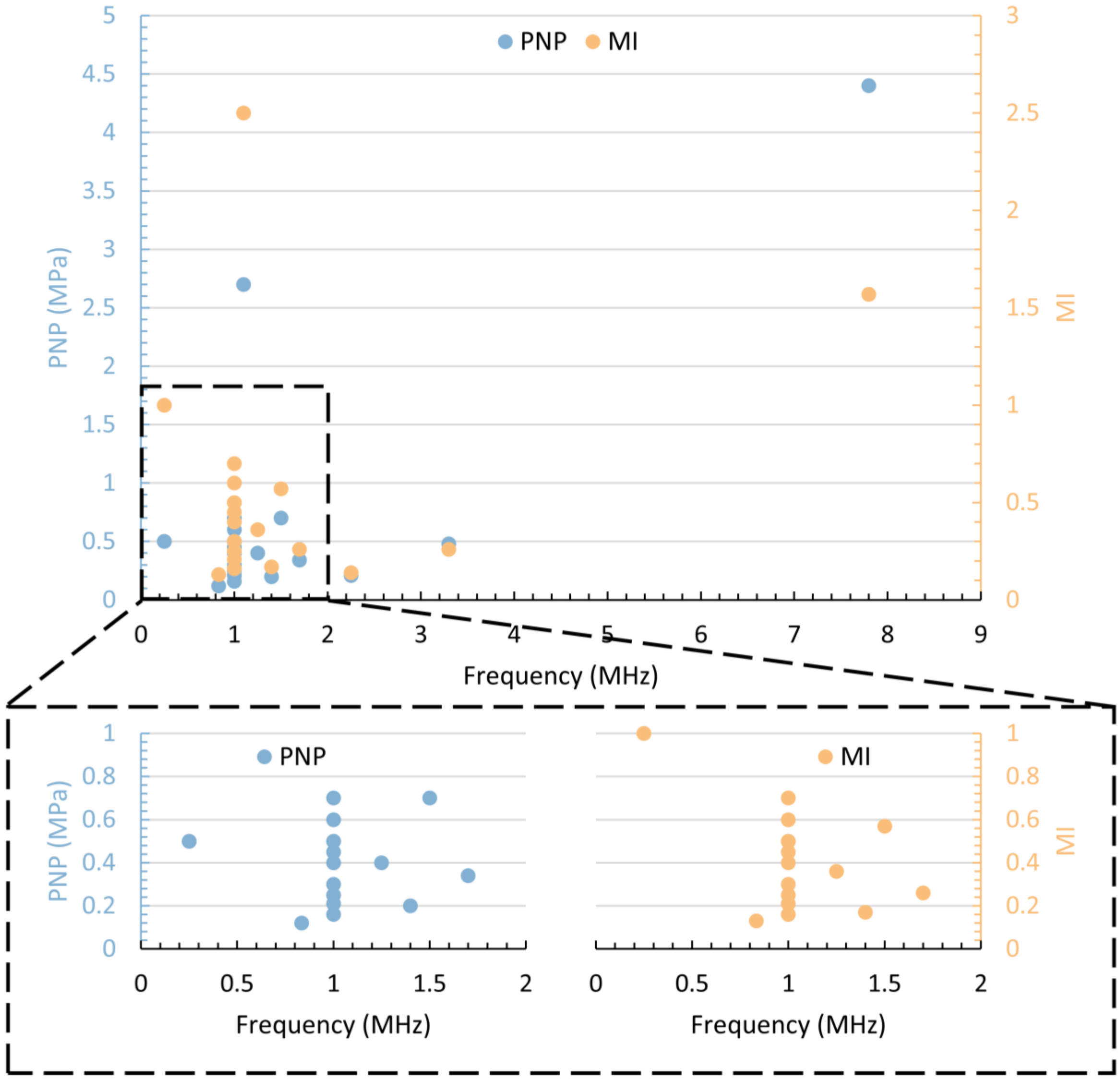


**Fig. S3.** Summary of appropriate sonication parameters (including frequency, PNP, and MI) for *in-vitro* sonoporation cell tests.

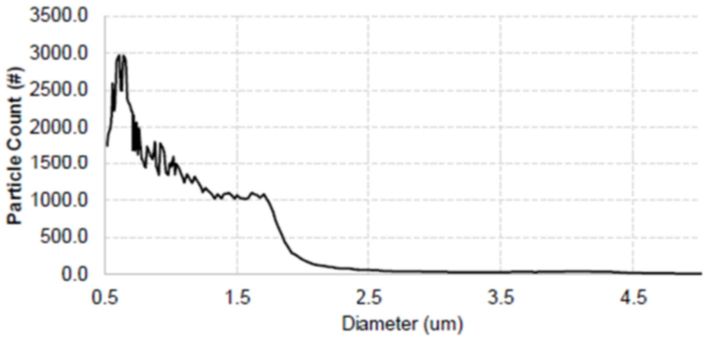


**Fig. S4.** Microbubble size distribution provided by the vendor (SonoVol Inc., USA).

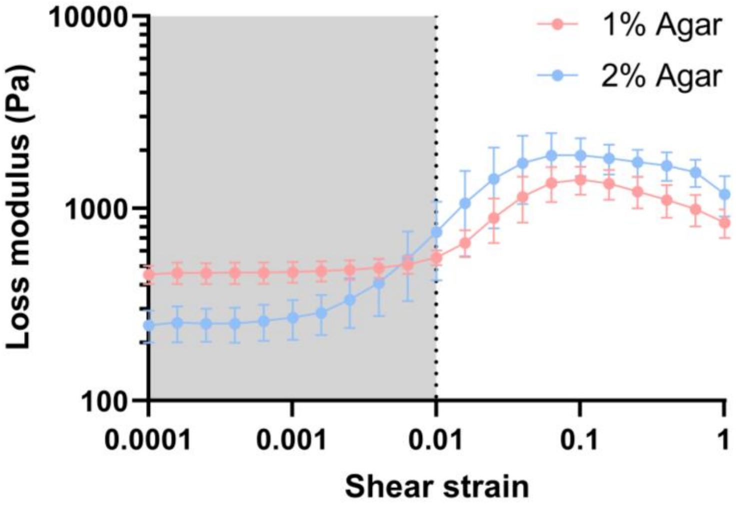


**Fig. S5.** Measured shear loss modulus (G") of 1% and 2% agar phantom (n = 3).

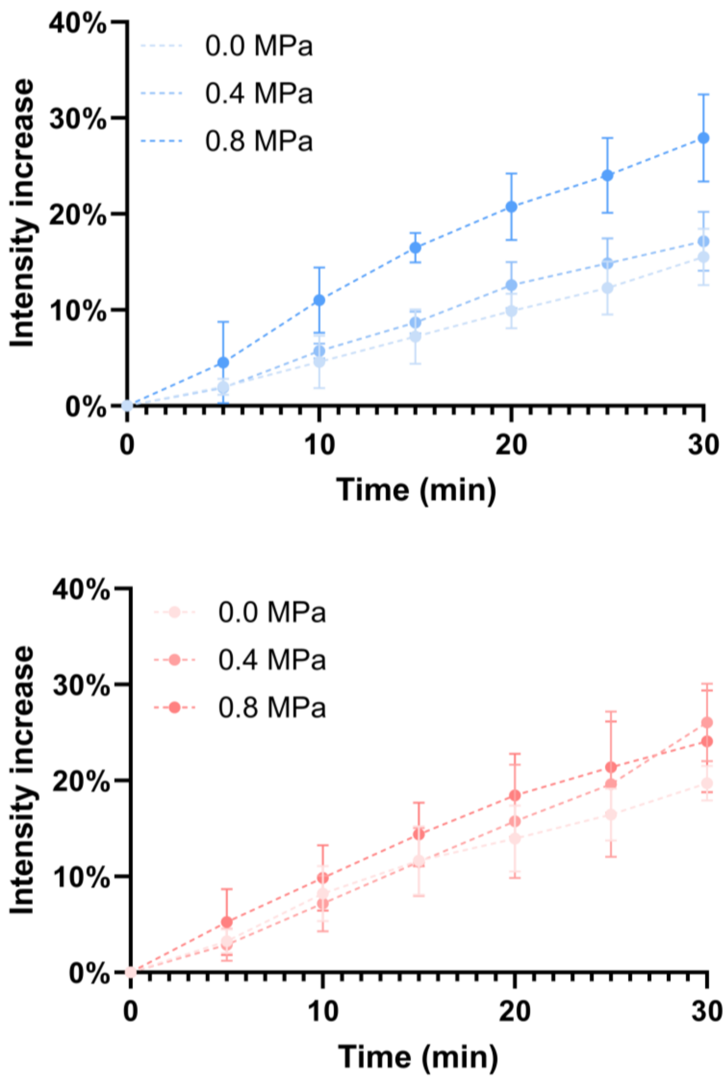


**Fig. S6.** Measured intensity changes of the dye diffusion area for demonstrating drug penetration along the y-axis in 1% (top) and 2% (bottom) agar phantom (n = 3).

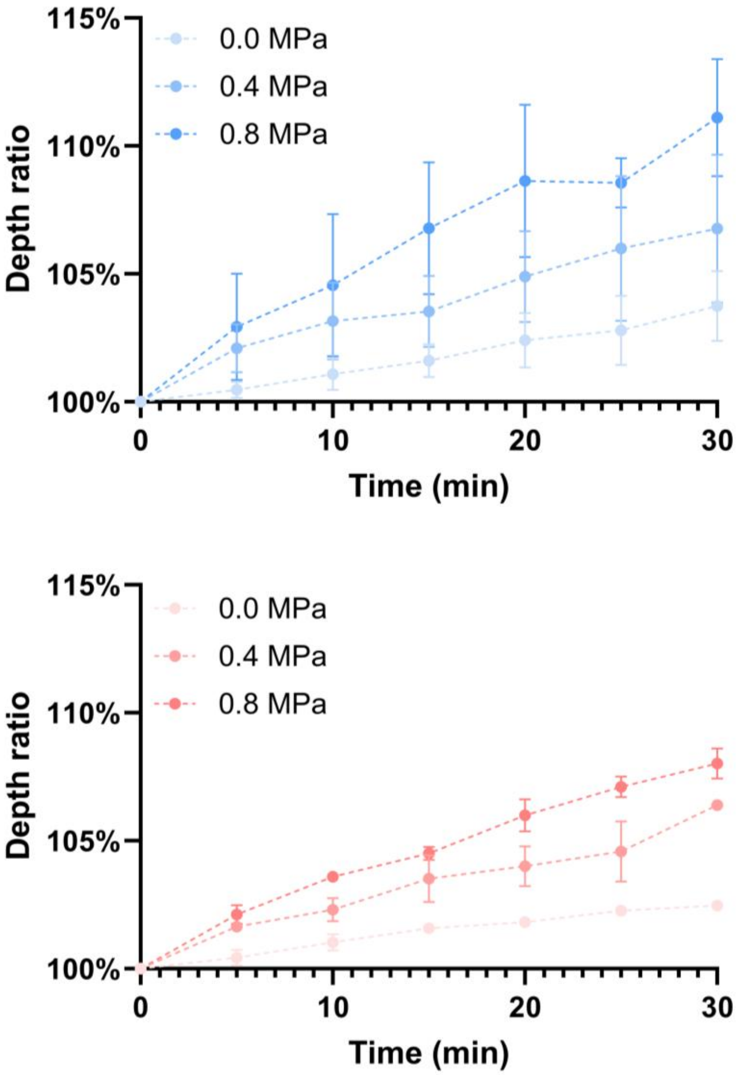


**Fig. S7.** Measured depth changes of the dye diffusion area for demonstrating drug penetration along the z-axis in 1% (top) and 2% (bottom) agar phantom (n = 3).

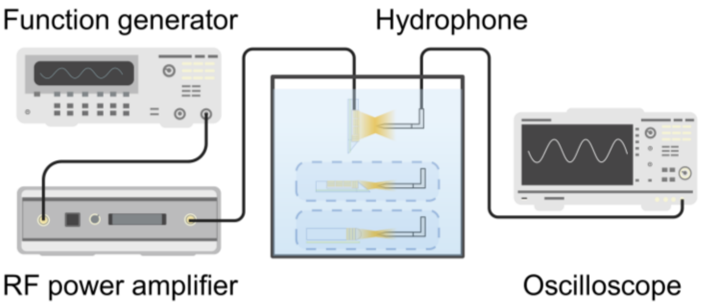


**Fig. S8.** Experiment setup for measuring the acoustic output from cPULSE and nPULSE (sideward and forward direction).
